# Short OGA is targeted to the mitochondria and regulates mitochondrial reactive oxygen species level

**DOI:** 10.1101/2021.12.25.474160

**Authors:** Patrick Pagesy, Abdelouhab Bouaboud, Zhihao Feng, Philippe Hulin, Tarik Issad

## Abstract

O-GlcNAcylation is a reversible post-translational modification involved the regulation of cytosolic, nuclear and mitochondrial proteins. Only two enzymes, OGT and OGA, control attachment and removal of O-GlcNAc on proteins, respectively. Whereas a variant OGT (mOGT) has been proposed as the main isoform that O-GlcNAcylates proteins in mitochondria, identification of a mitochondrial OGA has not been performed yet. Two splice variants of OGA (short and long isoforms) have been described previously. In this work, using cell fractionation experiments, we show that short-OGA is preferentially recovered in mitochondria-enriched fractions from HEK-293T cells as well as mouse embryonic fibroblasts. Moreover, fluorescent microscopy imaging confirmed that GFP-tagged short-OGA is addressed to mitochondria. In addition, using a BRET-based mitochondrial O-GlcNAcylation biosensor, we show that co-transfection of short-OGA markedly reduced O-GlcNAcylation of the biosensor, whereas long-OGA had no significant effect. Finally, using genetically encoded or chemical fluorescent mitochondrial probes, we showed that short-OGA overexpression increases mitochondrial ROS levels, whereas long-OGA had no significant effect. Together, our work reveals that the short-OGA isoform is targeted to the mitochondria where it regulates ROS homoeostasis.

## Introduction

O-GlcNAcylation is a post-translational modification corresponding to the attachment of a single O-linked N-acetylglucosamine (O-GlcNAc) to serine or threonine residues of cytosolic, nuclear or mitochondrial proteins. This reversible modification regulates the localization, the activity and the stability of proteins according to the nutritional environment of the cell, and more specifically according to glucose availability. Only two enzymes regulate O-GlcNAc level on proteins: OGT (O-linked N-Acetylglucosamine transferase), a glycosyl transferase that adds O-GlcNAc to proteins, and OGA, a β-N-Acetylglucosaminidase, distinct from acidic lysosomal hexosaminidase, which removes the O-GlcNAc from proteins. Numerous studies indicated a dynamic cross-talk between O-GlcNAcylation and phosphorylation, permitting fine-tuning of cell signalling pathways and regulation of gene expression (Issad & Kuo, 2008; Issad *et al*, 2010). O-GlcNAcylation has been involved in important human pathologies, including neurogenerative diseases, diabetes and cancer (Bond & Hanover, 2013).

Whereas protein O-GlcNAcylation in the cytosol and nucleus has been largely investigated, relatively little is known about O-GlcNAc cycling enzymes and their targets in the mitochondria. Alternative splicing of OGT results in the production of 3 different mRNA isoforms (Hanover *et al*, 2003; Love *et al*, 2003; Sacoman *et al*, 2017; Trapannone *et al*, 2016), that can code for 3 different proteins: a nucleo-cytoplasmic long form (ncOGT), a short isoform (sOGT) also found in the cytosol and nucleus, and at least in humans and non-humans primates, a mitochondria-targeted variant (mOGT) (Supplementary figure 1). Several studies have shown the important role of O-GlcNAcylation in mitochondrial functions. Increases in O-GlcNAcylation of mitochondrial proteins have been observed upon high-glucose conditions (Hu *et al*, 2009; Makino *et al*, 2011), and recent work pointed to perturbation in the localisation of mOGT in cardiomyocytes mitochondria from diabetic mice (Banerjee *et al*, 2015). Several lines of evidence indicate that alteration of O-GlcNAc cycle disrupts mitochondrial homoeostasis (Akinbiyi *et al*, 2021; Jozwiak *et al*, 2021; Ma *et al*, 2015 ; Shin *et al*, 2011; Tan *et al*, 2017; Tan *et al*, 2014), including alteration in reactive oxygen species production (Jozwiak *et al*., 2021; Ngoh *et al*, 2011; Tan *et al*., 2017; Wang *et al*, 2016; Zhao *et al*, 2014).

Although OGA enzymatic activity has been demonstrated in mitochondria (Banerjee *et al*., 2015; Hu *et al*., 2009), the protein isoform involved has not been characterized. Alternative splicing of OGA also results in the production of two different mRNAs, coding for either long and short OGA isoforms (Kim *et al*, 2006). The long OGA comprise an O-GlcNAcase activity in its N-terminal side and pseudo histone acetyltransferase (HAT) domain in its C-terminal side (Hanover *et al*, 2012). In the human short OGA, the HAT domain is deleted, and a small intronic derived-sequence give rise to a unique 15 amino acids C-terminal peptide. While the long OGA (L-OGA) isoform has been largely studied and was shown to be mainly cytoplasmic and nuclear, conflicting results have been reported concerning the short (S-OGA) isoform. Indeed, in glioblastoma cells, Comtesse et al. detected by western-blot a band of 130 kDa corresponding to L-OGA in the cytosolic fraction, and a band of 75 kDa in the nuclear fraction that they assumed to be S-OGA (Comtesse *et al*, 2001). In contrast, by fluorescence microscopy, Hanover’s group did not detect GFP-tagged S-OGA in the nucleus but rather suggested a lipid-droplet localization for this protein in HeLa cells (Keembiyehetty *et al*, 2011).

Previous studies in cultured cells, animal tissues and human samples, have described a strong correlation between ncOGT and L-OGA expression (Kazemi *et al*, 2010; Pagesy *et al*, 2018; Slawson *et al*, 2005; Zhang *et al*, 2014), permitting tight control of O-GlcNAc level in the cell. While analysing OGT and OGA mRNA expression levels in human leukocytes from healthy donors, we noticed that in contrast to L-OGA mRNA which correlated with ncOGT mRNA, S-OGA mRNA expression did not correlate with ncOGT, but was tightly correlated with the mitochondrial mOGT mRNA (**Supplementary Figure 1**). This suggested a role for the short OGA isoform in the mitochondria and prompted us to evaluate its addressing to this organelle. We discovered that S-OGA is indeed the main OGA isoform in mitochondria and that it is involved in the control of ROS levels in this organelle.

## Results and discussion

Long OGA has a theoretical molecular weight of 102 kDa but is well known to run on SDS-PAGE with an apparent molecular weight of 130 kDa (Gao *et al*, 2001). To evaluate the migration profile of short OGA on SDS-PAGE, we first transfected HEK-293T cells with pcDNA3 or with plasmids coding for either S-OGA or L-OGA. Proteins from total cell lysates were submitted to SDS-PAGE followed by western-blotting using an antibody directed against a region common to both isoforms (residues 500-550, Novus Biologicals antibody). As shown in **Supplementary Figure 2A**, transfected S-OGA and L-OGA were readily detected in total cell lysate. As expected, L-OGA has an apparent molecular weight of about 130 kDa on SDS-PAGE. S-OGA, which has a theoretical molecular weight of 76 kDa, ran with an apparent molecular weight of about 95-100 kDa. Therefore, like L-OGA, S-OGA run on SDS-PAGE with an apparent molecular weight higher than predicted from its amino-acid sequence.

We then evaluated the relative distribution of endogenous L-OGA and S-OGA in total cell lysate (TCL), cytosolic (Cyto) and mitochondrial enriched (Mito) fractions from HEK-293-T cells. In total cell lysates, we detected a major band of about 130 kDa corresponding to the expected molecular weight of the long OGA, and fainter bands, including a band of about 95 kDa, possibly correponding to the short OGA isoform **(Fig. 1A)**. Cell fractionation experiments indicated that L-OGA was mainly recovered in the cytosolic fraction, whereas S-OGA was essentially recovered in the mitochondrial enriched fraction, although a band corresponding to L-OGA and an additional band of higher molecular weight were also detected in this fraction. Densitometric analysis of the blots indicated that the relative amount of S-OGA over L-OGA was much higher in the mitochondrial-enriched fraction **(Fig. 1A and B)**.

**Figure 1:**
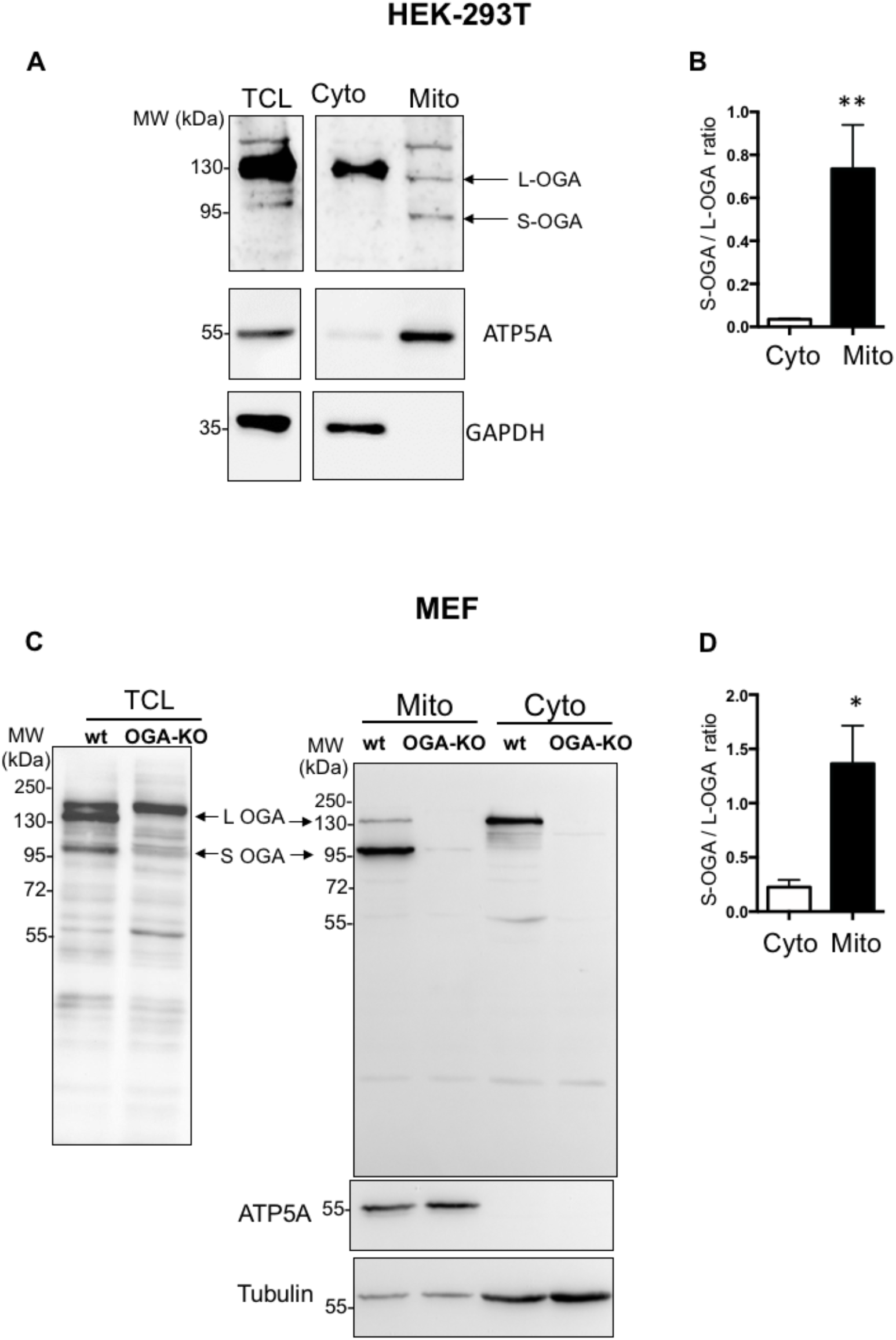
Detection of long and short OGA isoforms in HEK-293T cells and mouse embryonic fibroblasts (MEF). (A) Detection of endogenous S-OGA (95 kDa) and L-OGA (130 kDa) bands in total cell lysates (TCL), mitochondrial enriched (Mito) and cytosolic enriched (Cyto) fractions from HEK-293T cells by western-blotting with anti-OGA Novus antibody. Mitochondrial and cytosol fractions were controlled using anti-ATP5A and anti-GAPDH antibodies. The 95 kDa band was essentially recovered in the mitochondria-enriched fraction, whereas the 130 kDA band was mostly recovered in the cytosol-enriched fraction. (B) Densitometric analysis of the 95 kDa and 130 kDa OGA bands in HEK-293T cells. The results are the mean ± SEM of the ratio of the 95 kDa to the 130 kDa OGA bands detected in cytosol- and mitochondria-enriched fractions (n=9; **: p<0.01). (C) Detection of endogenous S-OGA (95 kDa) and L-OGA (130 kDa) bands in total cell lysates (TCL), mitochondria-enriched (Mito) and cytosol-enriched (Cyto) fractions from wild-type (wt) and OGA-KO mouse embryonic fibroblasts (MEF) by western-blotting with anti-OGA Novus antibody. Mitochondrial and cytosol fractions were controlled using anti-ATP5A and anti-tubulin antibodies. The 95 kDa band was predominant in the mitochondria-enriched fraction, whereas the 130 kDa band was predominant in the cytosol-enriched fraction of wt-MEF. Both bands were absent in the corresponding fractions in OGA-KO MEF. An additional band, (about 180 kDa) was also detected in total cell lysates, but this band was present in both wt and OGA-KO MEF, indicating that it corresponded to an unrelated protein. (D) Densitometric analysis of the 95 kDa and 130 kDa OGA bands in wt-MEF. The results are the mean ± SEM of the ratio of the 95 kDa to 130 kDa OGA signals detected in cytosol- and mitochondria-enriched fractions (n=6; *: p<0.05).

Because antibodies quite often recognize non-specific bands that can be mistaken for the proteins of interest (Trapannone *et al*., 2016), we wanted to ensure that the 130 kDa and 95 kDa bands detected by Novus anti-OGA antibody indeed corresponded to OGA by using cytosol and mitochondria enriched fractions of embryonic fibroblasts (MEF) from wild-type and OGA-KO mice (St Amand *et al*, 2018) **(Fig. 1C)**. These two bands were detected in total cell lysates from wt-MEF, but not in OGA-KO MEF (left blot). An additional band of about 180 kDa was also detected in total cell lysates, but it was present in both wt and OGA-KO MEF, indicating that it corresponded to an unrelated protein recognized by this antibody. Cell fractionation of MEF showed that S-OGA was clearly the predominant form in the mitochondria-enriched fraction while L-OGA was predominant in the cytosol enriched fraction (**Fig. 1C and D**). These bands were undetectable in the corresponding fractions from OGA-KO MEF, confirming the identity of the bands and the good specificity of the antibody.

To further establish the cellular localization of S-OGA and L-OGA, we transfected GFP-tagged version of these proteins in HEK-293T cells (**Suppl. Figure 2B**). In agreement with Keembiyehetty et al. (Keembiyehetty *et al*., 2011), transfected S-OGA-GFP and L-OGA-GFP were detected with an anti-GFP antibody as bands of about 130 kDa and 160 kDa respectively. As shown in **Supplementary Figure 2B**, L-OGA-GFP was more abundant than S-OGA-GFP in the cytosolic enriched fraction. In contrast, S-OGA-GFP was more abundant than L-OGA-GFP in the mitochondria-enriched fraction, despite higher expression of L-OGA-GFP in total cell lysates, confirming the preferential expression of S-OGA in the mitochondria (**Suppl. Fig. 2B, C**). Similar results were obtained in MEF transfected with these constructs (**Suppl. Fig. 2D, E**), Thus, although S-OGA is clearly preferentially recovered in the mitochondrial-enriched fractions, both S-OGA and L-OGA isoforms were found in both compartments. Nonetheless, mitochondrial and cytosol-enriched fractions may have cross-contaminated each other during the centrifugation procedure, resulting in the recovery of S-OGA in the cytosolic fraction and L-OGA in the mitochondrial fraction. To confirm specific localization of S-OGA in the mitochondria, we used confocal microscopy. Since no antibodies are available to specifically detect the short OGA isoform by cell imaging, we co-transfected HEK-293T cells with cDNAs coding for either GFP-tagged short or long OGA isoforms and a mitochondrial-targeted mCherry protein. Confocal microscopy imaging (**Fig. 2**) revealed that L-OGA isoform was mainly found in the cytosol and the nucleus (**Fig. 2A**), whereas S-OGA was not detected in the nucleus (**Fig. 2B**), in agreement with Keembiyehetty et al observation (Keembiyehetty *et al*., 2011). In contrast, S-OGA isoform co-localized with mitochondrial mCherry protein (**Fig. 2B**), indicating specific mitochondrial addressing of this isoform. Mitochondrial localisation of the short OGA isoform was confirmed using structured illumination microscopy (Turkowyd *et al*, 2016)(**Fig. 2C and D**).

**Figure 2:**
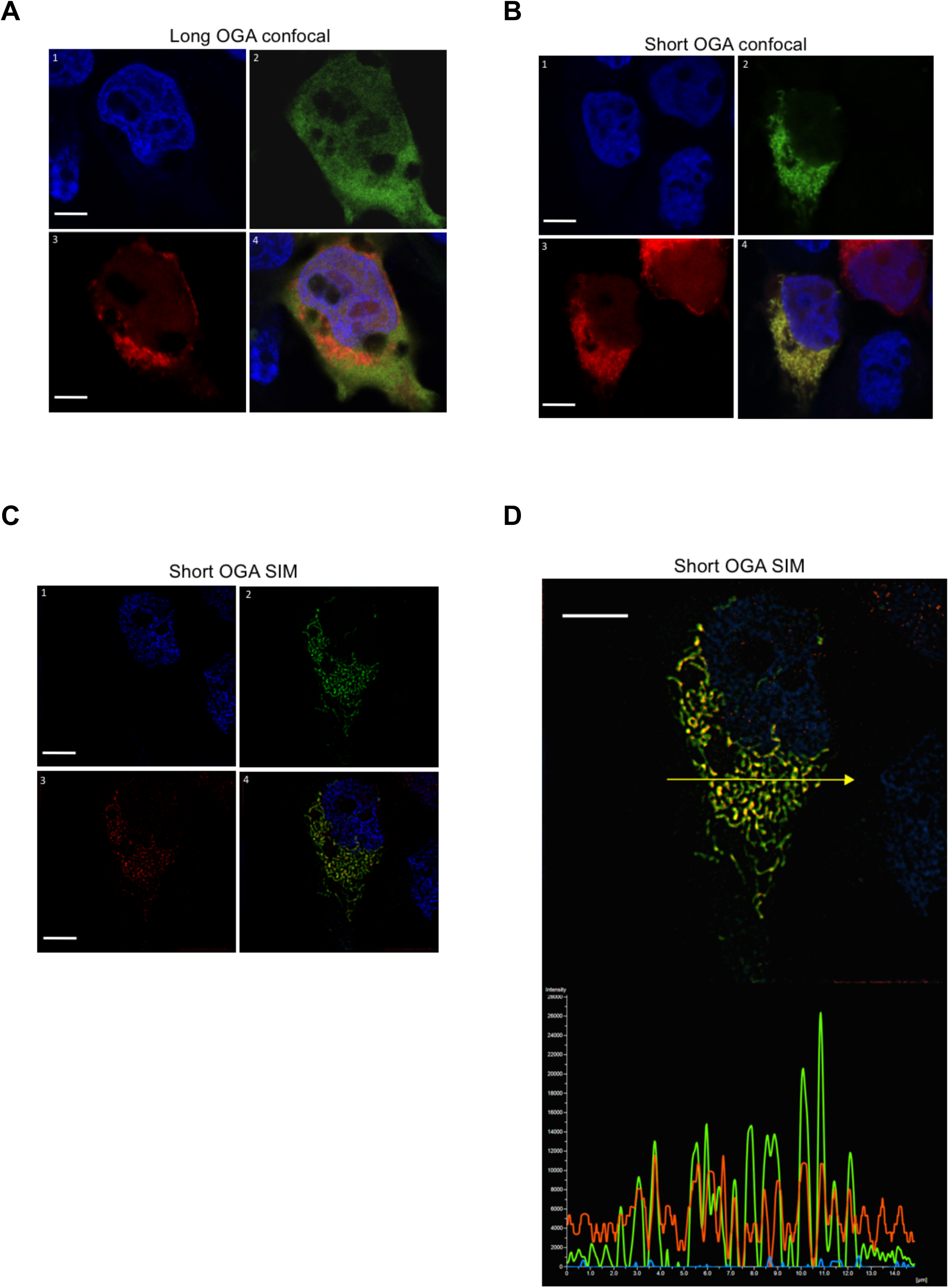
Detection of GFP-tagged long and short OGA isoforms in HEK-293T cells. HEK-293T cells were transfected with mitochondrial-targeted mCherry and either GFP-tagged short or long OGA isoforms, fixed, stained with DAPI for visualisation of the nuclei (blue, panels 1), and analysed by confocal microscopy in the GFP (green, panels 2) and mCherry (red, panels 3) channels. Merge images are shown in panels 4. (A) the long OGA isoform was detected in the cytosol and the nucleus but poorly co-localized with mito-cherry labelled mitochondria. (B) Co-localization of the short OGA isoform with mito-mCherry in a cell expressing GFP-tagged short OGA. (C, D) Structured illumination microscopy (SIM) confirmed localisation of S-OGA in the mitochondria. (C) Separated SIM and merge images of this cell in the GFP, mCherry and Dapi channels are shown. (D) Densitometric analysis of the different signals detected along the arrow shown on the upper panel indicates co-localisation of S-OGA-GFP with mito-mCherry (green: S-OGA-GFP, red: mito-mCherry, blue: Dapi staining of the nucleus, white bar scale: 5µm). Representative images are shown.

S-OGA results from an mRNA splicing that retains an intronic sequence coding for a 15 amino-acid peptide located in the C-terminus of the human short OGA isoform. This peptide sequence, which is not present in L-OGA, is fully conserved in five primate species and partially conserved in other mammalian species (**Suppl. Fig.3**). We hypothesized that this sequence may serve as a mitochondria-targeting sequence. To test this hypothesis, we introduced the sequence coding for this Intron-derived Peptide (IdP) at the C-terminus of the GFP. Transfection of HEK-293T cells with cDNA coding for either GFP or GFP-IdP showed that GFP-IdP was essentially recovered in the mitochondria enriched fraction, in contrast to GFP alone which was essentially recovered in the cytosolic fraction (**Fig. 3A and B)**. Confocal microscopy experiments confirmed mitochondrial localisation of IdP-GFP in HEK-293T as well as in HeLa cells (**Fig. 3C)**. This result suggests that the intron-derived peptide may acts as a mitochondria-addressing sequence for S-OGA.

**Figure 3:**
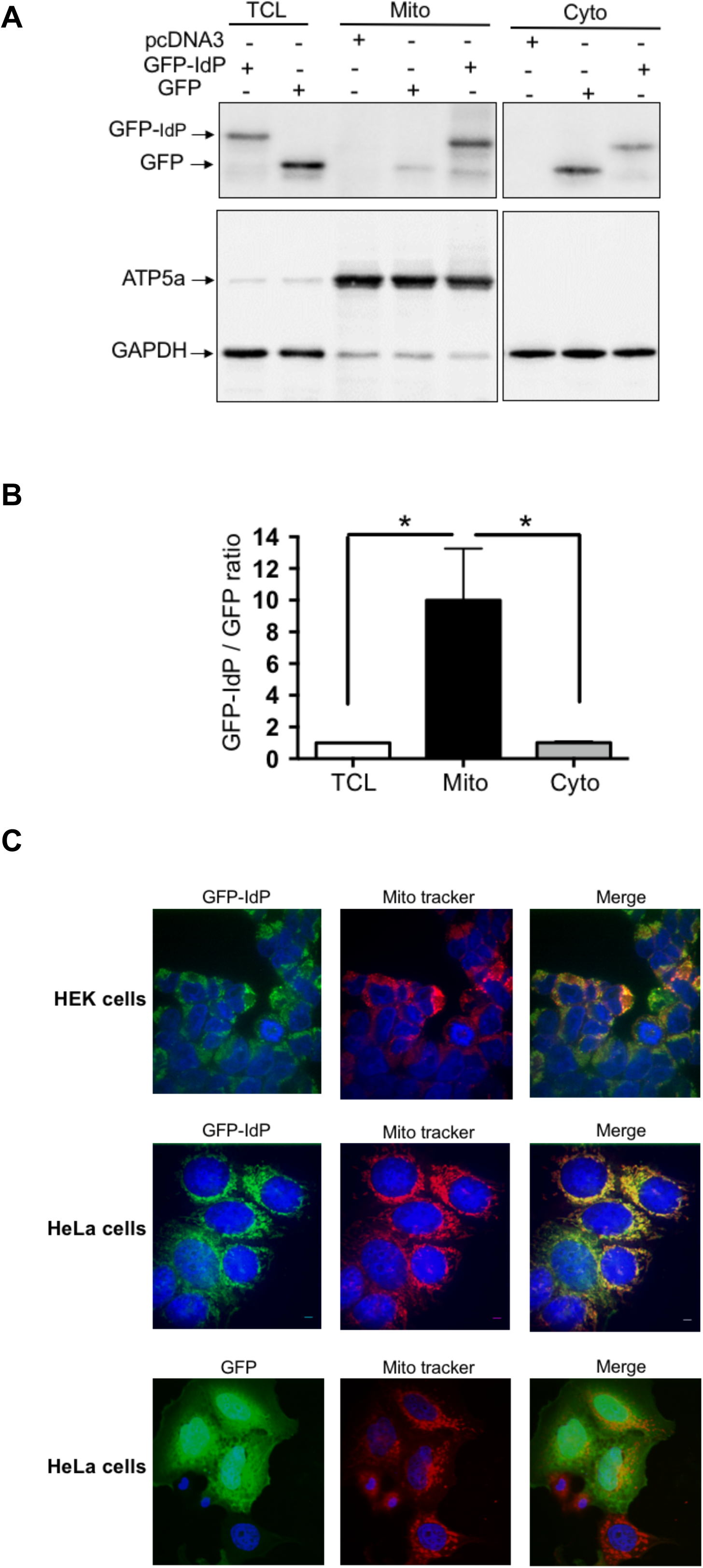
The intron-derived peptide of short OGA is sufficient to target GFP to the mitochondria. (A) HEK-293T cells were transfected with cDNA coding for either GFP or GFP fused at its C-terminus with OGA intron-derived peptide (GFP-IdP). Total cell lysate, mitochondrial and cytosol enriched fractions were analysed by western-blotting with anti-GFP antibody. GFP-IdP was more abundant in the mitochondrial-enriched fraction whereas GFP was more abundant in the cytosol enriched fraction. (B) Densitometric analysis of the GFP and GFP-IdP bands. The results are the mean ± SEM of the ratio of the GFP-IdP to GFP signals detected in cytosol- and mitochondria-enriched fractions (n=5; *: p<0.05). (C) HEK-293T cells and HeLa cells were transfected with cDNA coding GFP fused at its C-terminus with OGA intron-derived peptide (GFP-IdP), fixed and labelled with mitotracker. Confocal microscopy experiments confirmed co-localisation of GFP-IdP with mitotracker. GFP alone was widely distributed into the cells, including the nucleus (HeLa cells).

To determine whether S-OGA expression can regulate protein O-GlcNAcylation in mitochondria, we used a BRET-based O-GlcNAc biosensor that monitors O-GlcNAcylation activity in living cells (Al-Mukh *et al*, 2020; Groussaud *et al*, 2017). This biosensor comprises a lectin (GafD) domain fused to a luciferase and an O-GlcNAcylation substrate peptide derived from casein kinase 2 fused to a Venus fluorescent protein (**Supplementary Fig. 4**). O-GlcNAcylation of CKII promotes its binding to GafD, resulting in an increased in BRET signal (Al-Mukh *et al*., 2020), whereas removal of the GlcNAc by OGA will decrease BRET signal (Groussaud *et al*., 2017) (**Suppl. Fig. 4**). To specifically monitor O-GlcNAcylation in mitochondria, the mitochondrial targeting sequence of cytochrome oxidase subunit 8A (COX8A) was fused the cDNA coding for the BRET-O-GlcNAc biosensor (Mitochondrial O-GlcNAc BRET biosensor, **Suppl. Fig. 4**). HEK293-T cells were co-transfected with this biosensor and either pcDNA3, S-OGA or L-OGA. We observed that co-transfection of S-OGA markedly reduced basal BRET, whereas L-OGA had no significant effect on BRET signal (**Suppl. Fig. 4 and Fig. 4A**). These results strongly suggest that S-OGA is indeed an important regulator of protein O-GlcNAcylation level in the mitochondria.

**Figure 4:**
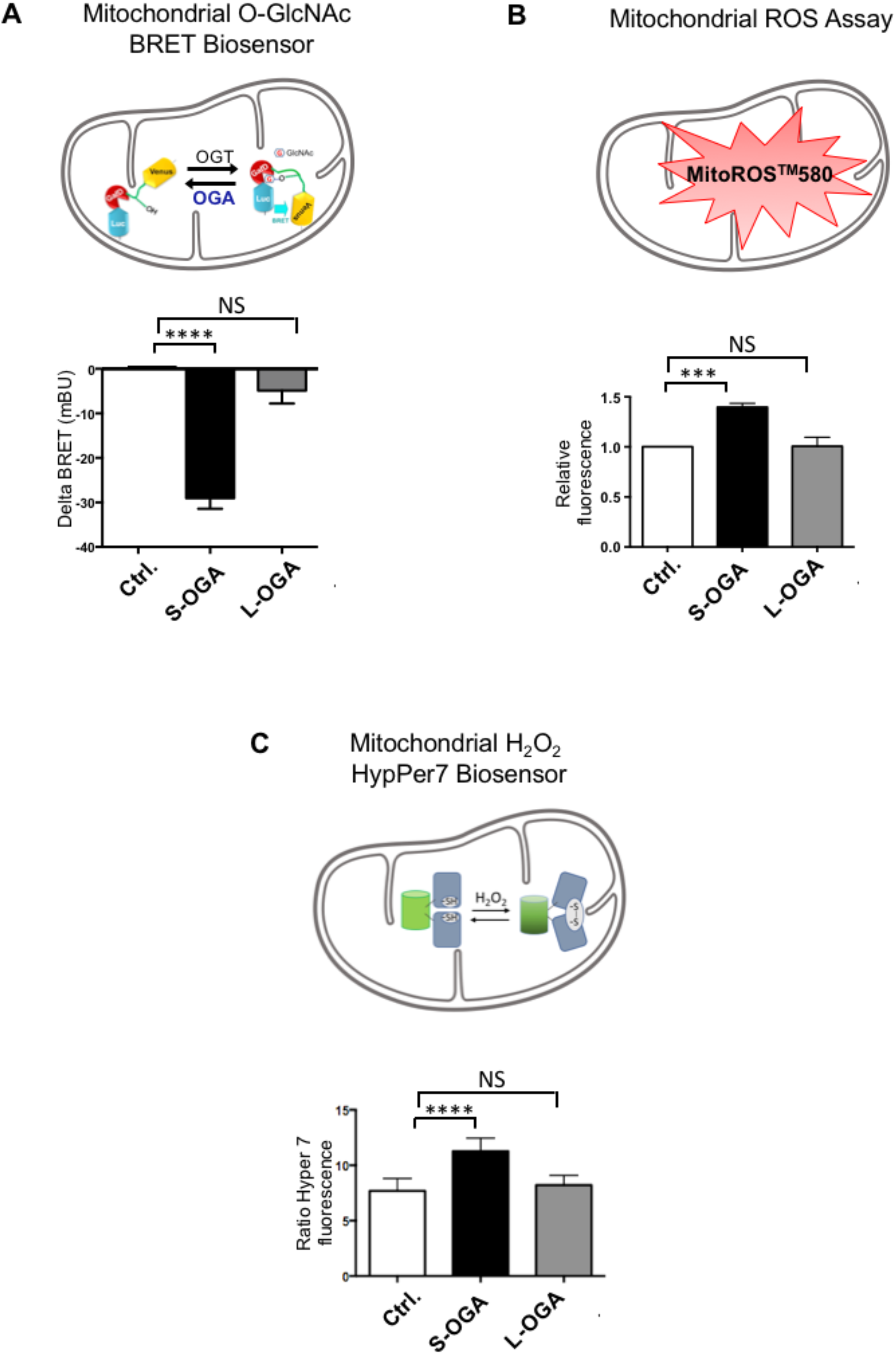
Short OGA expression reduced O-GlcNAcylation and increase ROS levels in the mitochondria. (A) HEK-293T cells were co-transfected with the mitochondria-targeted BRET biosensor and either pcDNA3, S-OGA or L-OGA. The graph shows the difference in BRET signal (delta BRET, expressed in milliBRET Units) between pcDNA3 transfected cells and either S-OGA or L-OGA transfected cells. Data are the mean ± SEM of 9 independent BRET experiments. Statistical analysis was performed using ANOVA followed by Dunnett’s post-test (****: p<0.0001, when compared to the control condition, NS: non-significative). S-OGA markedly reduced BRET signal, indicating decreased O-GlcNAcylation of the mitochondrial O-GlcNAc biosensor by this isoform. (B) HEK-293T cells transfected with pcDNA3, S-OGA or L-OGA were incubated for 1h with MitoROS™580. Fluorescence emission at 590 nm was then measured after excitation at 540 nm. Results are expressed as fluorescence in S-OGA or L-OGA transfected cells relative to pcDNA3 transfected cells, and are the mean ± SEM of 5 independent experiments. (C) HEK-293T cells were co-transfected with the mitochondria H_2_O_2_ biosensor (HyPer7) and either pcDNA3, S-OGA or L-OGA. In each experiment, the ratio of fluorescence emission at 516 nm after excitation at 499 nm to fluorescence emission at 516 nm after excitation at 400 nm was determined. Results are expressed as fold-change in 485/390 ratio in S-OGA or L-OGA transfected cells compared to pcDNA3 transfected cells, and are the mean ± SEM of 13 independent experiments.

Several studies have indicated that O-GlcNAcylation is involved in the regulation of oxidative stress (Chen *et al*, 2018). Since mitochondria is an important player in ROS production, we evaluated the effect of S-OGA overexpression on mitochondrial ROS levels in HEK-293T cells. Using a mitochondrial superoxide detection assay (MitoROS™580), we observed that S-OGA overexpression in HEK-293T cells significantly increased mitochondrial ROS level, when compared to pcDNA transfected cells, whereas L-OGA had no significant effect (**Fig. 4B**).

In the mitochondria, ROS are readily converted into more stable and less toxic hydrogen peroxide H^2^O_2_ by superoxide dismutase (Candas & Li, 2014). HEK-293T cells were co-transfected with pcDNA3, S-OGA or L-OGA, and a mitochondrial GFP-derived H_2_O_2_ biosensor (HyPer7, (Pak *et al*, 2020)), which permits to specifically monitor H_2_O_2_ level in the mitochondria. We observed that S-OGA expression resulted in a significant increase in H_2_O_2_ in the mitochondria, whereas L-OGA had no significant effect (**Fig. 4C**).

Together, our results indicate that S-OGA in the mitochondria regulates protein O-GlcNAcylation, superoxide and hydrogen peroxide levels in mitochondria.

## Conclusion

In this work, we discovered that S-OGA is preferentially targeted to the mitochondria, where it appears to modulate ROS levels. Mitochondria is believed to be the main source of cellular ROS. Whereas H_2_O_2_ can act as a signalling molecule in the cell, excess ROS production and elevated H_2_O_2_ levels have deleterious effects and are involved in several pathological conditions, including cancer, inflammatory diseases, type 2 diabetes, neurodegenerative diseases, and aging. The use of mitochondria-targeted small molecules has been proposed as a potential therapeutic strategy, most notably to specifically deliver antioxidants to this compartment (Oliver & Reddy, 2019; Smith *et al*, 2011). The discovery that S-OGA is addressed to the mitochondria, and that it plays a role in the regulation of ROS levels in this organelle, may open new avenues for the development of molecules with potential therapeutic value. To this aim, it will be necessary to fully characterize S-OGA targets in the mitochondria, to elucidate its mechanism of action in the regulation of ROS production, and to develop molecules that will specifically target S-OGA activity and/or interaction with its mitochondrial protein partners.

## Material and Methods

### Antibodies

Anti-OGA antibody (NBP2-32233) was from Novus Biologicals. Anti-ATP-5A antibody (15H4C4) was from Abcam. Anti-GFP antibody was from Roche (clones 7.1-13.1). Anti-human GAPDH (sc-47724) and anti-alpha Tubulin (sc-8035) antibodies were from Santa Cruz.

### Expression of S- and L-OGA mRNA in human leukocytes

Human leukocytes were obtained from blood samples of healthy volunteers (age 44.7 ± 1.7, 32 females, 35 males) from the French blood Agency (Etablissement Français du Sang, Ile-de-France, Site Trinité; Agreement number INSERM-EFS:18/EFS/030). For each individual, 5-10 ml of blood were collected in EDTA tubes. Leucocytes were isolated after red blood cell lysis in 3 volumes of RBC Lysis Buffer (Santa Cruz). Leucocytes were pelleted by centrifugation at 280 g during 5 min. This procedure was repeated once or twice to eliminate residual red blood cells. The pellet was then washed in PBS, lysed in Trizol, and total RNA were isolated as described previously (Strobel *et al*, 1999). RT-qPCR were performed as described previously (Pagesy *et al*., 2018) using the primers indicated in Supplementary Table 1.

### Preparation of cytosolic and mitochondrial-enriched fractions

HEK-293T cells were cultured as described previously (Blanquart *et al*, 2006). Preparation of mitochondrial-enriched fractions from HEK-293T cells was performed as described below. Briefly, HEK-293T cells were cultured to confluence in T-175 flasks. Cells from one T-175 flask were collected by trypsin digestion, washed in PBS and re-suspended in 1ml of ice-cold homogenization buffer containing 10 mM Hepes, pH 7.4, 250 mM sucrose, 1 mM AEBSF and 1mg/ml of pepstatin, antipain, leupeptin and aprotinin. Cells were then homogenized on ice by flushes (30 up and down) through a 1ml syringue with a 22G needle. Homogenates were then centrifuged twice at 1000 g for 10 min to pellet and discard nuclei and large debris. Mitochondria were then pelleted by centrifugation at 11000 g during 10 min, washed with 1ml homogenization buffer, centrifuged at 11000 g for 10 min and resuspend in the same buffer. The supernatant of the first 11000 g centrifugation, mainly containing the cytosolic fraction, and the mitochondrial-enriched pellet fraction, were then stored at -80°C for subsequent analysis by western-blotting.

15-45 µg of proteins from either cytosolic or mitochondrial enriched fractions were submitted to western-blotting as described previously (Liu *et al*, 1998). Mitochondrial and cytosolic fractions were controlled using anti-ATP5A and anti-human GAPDH or anti-alpha tubulin antibodies.

### Confocal microscopy experiments

HEK-293T cells plated on polylysine-coated coverslips were transfected with cDNA (100 ng/40000 cells) coding for mitochondrial-targeted mCherry (mCherry-Mito-7, a gift from Michael Davidson (Addgene plasmid # 55102), (Olenych *et al*, 2007)) and either GFP-tagged long or short OGA isoforms (gifts from John A. Hanover). 48h after transfection, cells were fixed with 4% paraformaldehyde and stained with DAPI (4’,6-diamidino-2-phenylindole) for visualisation of the nuclei. Coverslips were sealed with ProLong diamond anti-fade mounting media (ThermoFisher Scientific) and analysed by confocal microscopy. Confocal and Structured Illumination Microscopy (SIM) was performed on the MicroPICell Facility of the University of Nantes using an inversed confocal Nikon A1 microscope coupled to the super resolution N-SIM. Z stack of 0.12 microm were performed using a 100× oil-immersion lens with high NA (SR ApoTIRF 100×, oil, NA: 1.49, Nikon). Images were acquired using NIS 4.2 software.

HeLa cells plated on coverslips in a 6 well plate (3×10^5^ cells/well) were transfected with 10 ng of cDNA coding for GFP or GFP-IdP using lipofectamine 2000 (Thermofisher Scientific). Cells were labelled with 200nM MitoTracker (Thermofisher Scientific) during 45 min at 37°C and then fixed with 4% paraformaldehyde, stained with DAPI (4’,6-diamidino-2-phenylindole) for visualisation of the nuclei and analysed by confocal microscopy using an inverted microscope (Leica DMI6000, objective lens 100×).

### BRET experiments

The coding sequence of mitochondrial targeting sequence from human COX8A was inserted upstream of the cDNA coding for the general O-GlcNAc-BRET biosensor described previously (Al-Mukh *et al*., 2020; Groussaud *et al*., 2017). HEK-293T cells were co-transfected with this mitochondrial O-GlcNAc-BRET biosensor and either pcDNA3, S-OGA or L-OGA plasmids. Cells transfected with luciferase alone and either pcDNA3, S-OGA or L-OGA plasmids were used to correct for background signal. BRET experiments were then performed exactly as described previously (Lacasa *et al*, 2005) using the Tristar2 LB 942 (Berthold) plate reader. Briefly, cells were pre-incubated for 5 min in PBS in the presence of 5µM coelenterazine. Each measurement corresponded to the signal emitted by the whole population of cells present in a well. BRET signal was expressed in milliBRET Unit (mBU). The BRET unit has been defined previously as the ratio 530 nm/485 nm obtained in cells expressing both luciferase and YFP, corrected by the ratio 530 nm/485 nm obtained under the same experimental conditions in cells expressing only luciferase (Issad *et al*, 2002; Nouaille *et al*, 2006). In each experiment, BRET signal was the mean of at least 10 successive measurement performed every min during at least 10-15 min. In each experiment, the mean of at least 10 repeated BRET measurements in a given experimental condition (see **supplementary Figure 4**) was taken as the BRET value obtained in this experimental condition (Al-Mukh *et al*., 2020). Delta BRET corresponded to the difference in BRET signal measured in cells transfected with pcDNA3 and cells transfected with either S-OGA or L-OGA.

### Determination of mitochondrial ROS using MitoROS probe

Mitochondrial ROS level was assessed using a Mitochondrial Superoxide Assay Kit according to the manufacturer instructions (Abcam). Briefly, HEK293-T cells (10^5^ cells/well in 12 well plates) were transfected with 500ng of pcDNA3 or cDNA coding for S-OGA or L-OGA using Lipofectamine 2000. 24h after transfection, cells were transferred into 96 well black microplate and cultured for an additional 24h. Cells were then incubated with the MitoROS™580 fluorescent probe (100 µl of MitoROS™580 stain working solution added in each well) for 1 h at 37°C. Fluorescence emission at 590 nm was then measured after stimulation at 540 nm using a CLARIOstar (BMG) fluorimeter.

### Determination of mitochondrial H2O2 using HyPer7 fluorescent probe

HyPer7 (pCS2+MLS-HyPer7, a gift from Vsevolod Belousov, Addgene plasmid #136470)) is an ultrasensitive fluorescent ratiometric probe for detection of mitochondrial H_2_O_2_. This GFP probe has two excitation maxima at 400 and 499 nm and one emission peak centred at 516 nm. Upon oxidation, excitation and absorption spectra of Hyper7 changes in a ratiometric way with a decrease at 400 nm and an increase of the 499 nm peak (Pak *et al*., 2020).

HEK293-T cells (10^5^ cells/well in 12 well plates) were co-transfected with 500ng of cDNA coding for HyPer7 and 500ng of pcDNA3 or cDNA coding for S-OGA or L-OGA using Lipofectamine 2000. 24h after transfection, cells were transferred into 96 well black microplate and cultured for an additional 24h. Culture medium was removed and the cells were washed with PBS and incubated in 100 µl PBS for fluorescence determination. Fluorescence was measured at 515/40 nm after excitation at 390/22 nm and 485/15 nm using a LB942 Tristar2 Berthold fluorometer. After removing background fluorescence, the ratio of fluorescence emission after excitation at 485 nm to fluorescence emission after excitation at 390 nm was taken as relative measurement of mitochondrial H_2_O_2_ levels. Results were expressed as fold-change in 485/390 ratio induced by S-OGA or L-OGA transfection compared to pcDNA3 transfected cells.

### Statistical analysis

Statistical analyses were performed using PRISM software. Comparisons between groups were performed using Student’s t test, or ANOVA followed by Dunnett’s post-test for multiple comparison analysis. Correlations were performed using Pearson analysis.

## Acknowledgments

We thank John A. Hanover for the plasmids coding for GFP-tagged short and long OGA isoforms, and for generously providing us with wt and OGA-KO mouse embryonic fibroblasts.

We acknowledge the IBISA MicroPICell facility (Biogenouest), member of the national infrastructure France-Bioimaging supported by the French national research agency (ANR-10-INBS-04) for confocal and SIM imaging of S-OGA-GFP.

## Authors contribution

PP, AB, ZF researched data, analyzed the results, contributed to discussion and edited the manuscript. PH performed SIM imaging. TI conceived the experiments, analyzed the data and wrote the manuscript.

## Figure legends

**Supplementary Figure 1:**
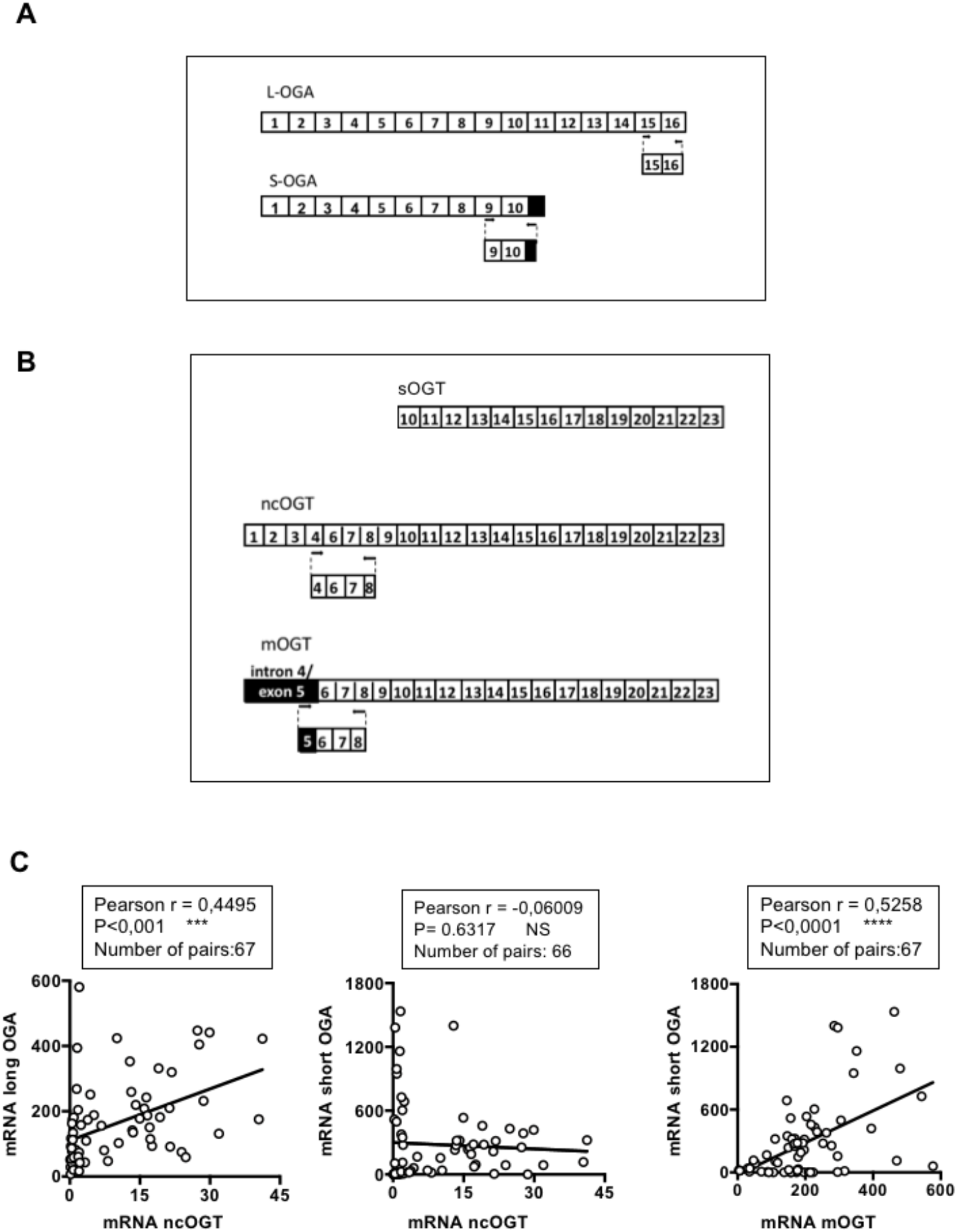
Correlations between S-OGA and mOGT mRNA expression in human leukocytes. (A) Alternative splicing of OGA results in the production of two different mRNAs, coding for either long or short OGA isoforms. The short mRNA isoform lacks the sequence coding for exons 11 to 16 but retains part of intron 10 as a coding sequence. The black box indicates the localization of the sequence that is specific for short OGA. 2 couples of primers were designed to specifically quantify mRNA expression of each isoform. (B) Alternative splicing of OGT results in the production of three different mRNAs, which can code for 3 different proteins: the two nucleo-cytoplasmic long (ncOGT) and short OGT variants (sOGT), and at least in human and other primates, a mitochondria-targeted variant (mOGT). The mOGT mRNA isoform is generated by the use of intron 4 as an alternative exon (exon 5). This transcript contains a unique ATG that produces a shorter isoform which comprises a 20 amino acid mitochondrial targeting sequence. 2 couples of primers were designed to specifically quantify mRNA expression of ncOGT and mOGT isoforms (the absence of any sequence specific to sOGT impairs evaluation of its expression). (C) Correlations between OGT and OGA mRNA splice variants were evaluated using Pearson’s analysis. OGA mRNA expression levels were measured in leucocytes from healthy donors by quantitative RT-PCR and normalized to the expression of cyclophilin A mRNA. Expression levels of ncOGT correlated with long OGA mRNA (left panel) but not with short OGA mRNA (middle panel), while mOGT mRNA expression levels correlated with short OGA (right panel).

**Supplementary Figure 2:**
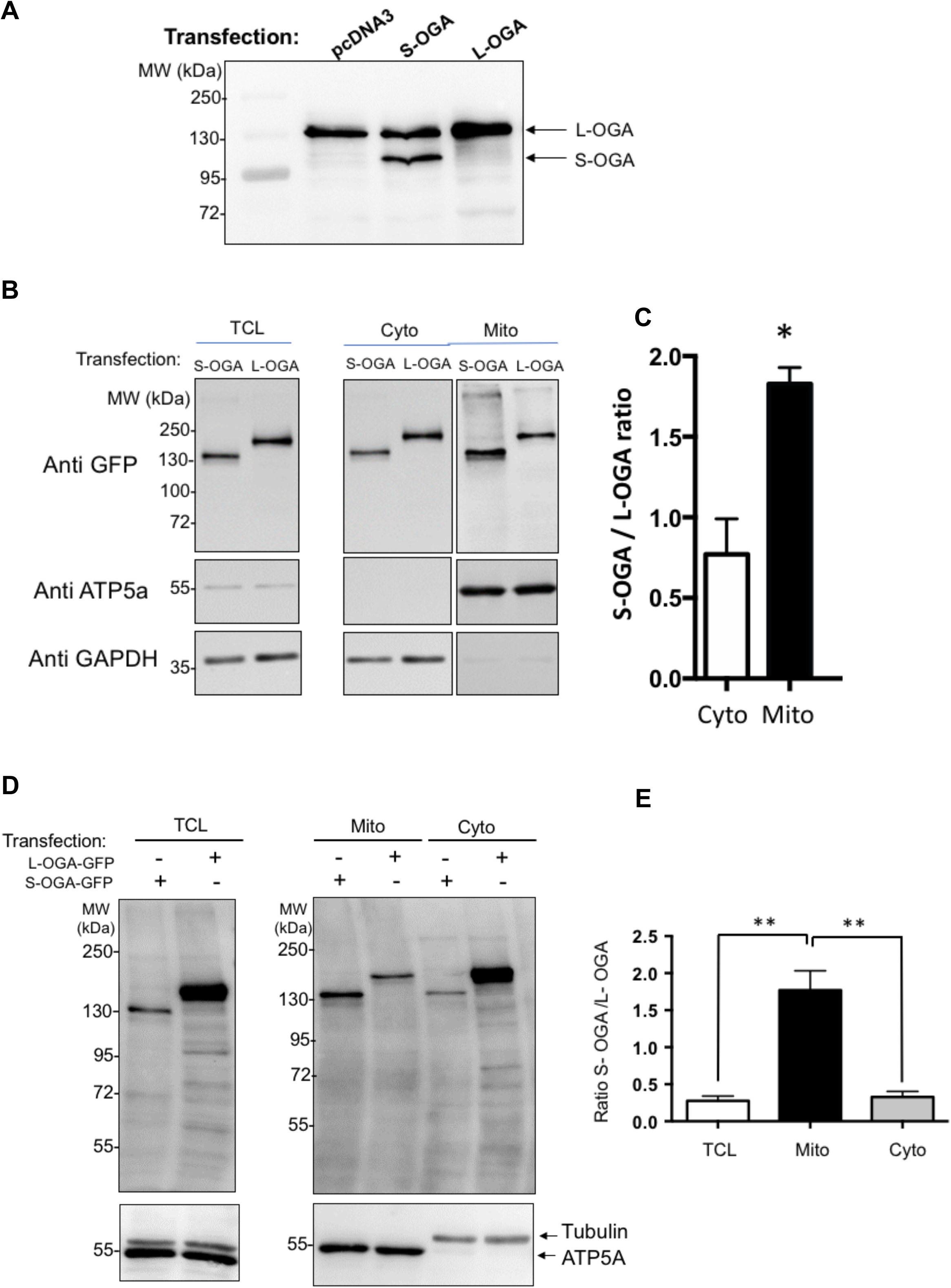
Expression of transfected S-OGA and L-OGA in HEK-293 T and mouse embryonic fibroblasts. (A) HEK-293T cells were transfected with pcDNA3 empty vector or plasmids coding for S-OGA or L-OGA. S-OGA and L-OGA in total cell lysates were detected as bands of apparent molecular weight of 95 kDa and 130 kDa by western-blotting using anti-OGA Novus antibody. (B) HEK-293T cells were transfected with plasmids coding for GFP-tagged S-OGA or L-OGA. GFP-S-OGA and GFP-L-OGA (left panel) were detected as bands of apparent molecular weights of 130 kDa and 160 kDa by western-blotting with an anti-GFP antibody. In mitochondria-enriched fractions from these cells, recovery of transfected GFP-S-OGA was higher than GFP-L-OGA, whereas GFP-L-OGA recovery was higher in the cytosolic enriched fraction. Mitochondrial and cytosol fractions were controlled using anti-ATP5A and anti-GAPDH antibodies. (C) Densitometric analysis of the 130 kDa and 160 kDa GFP-OGA bands in HEK-293T cells. The results are the mean ± SEM of the ratio of the 130 kDa to 160 kDa GFP-OGA bands detected in cytosol- and mitochondria-enriched fractions (n=3; *: p<0.05). (D) MEF were transfected with cDNA coding for either GFP alone, GFP-S-OGA (130 kDa) or GFP-L-OGA (160 kDa). (A) Total cell lysate (TCL), mitochondria (Mito) and cytosolic (Cyto) enriched fractions from these cells were submitted to western-blotting using an anti-GFP antibody. Mitochondrial and cytosol enrichment was controlled using anti-ATP5A and anti-α-tubulin antibodies. In mitochondria-enriched fractions, recovery of transfected GFP-S-OGA was higher than GFP-L-OGA, whereas GFP-L-OGA recovery was higher in the cytosolic enriched fraction. (E) Densitometric analysis of the 130 kDa and 160 kDa GFP-OGA bands in HEK-293T cells. The results are the mean ± SEM of the ratio of the 130 kDa to 160 kDa GFP-OGA signals detected in TCL, cytosol- and mitochondria-enriched fractions (n=3; *: p<0.05).

**Supplementary Figure 3.**
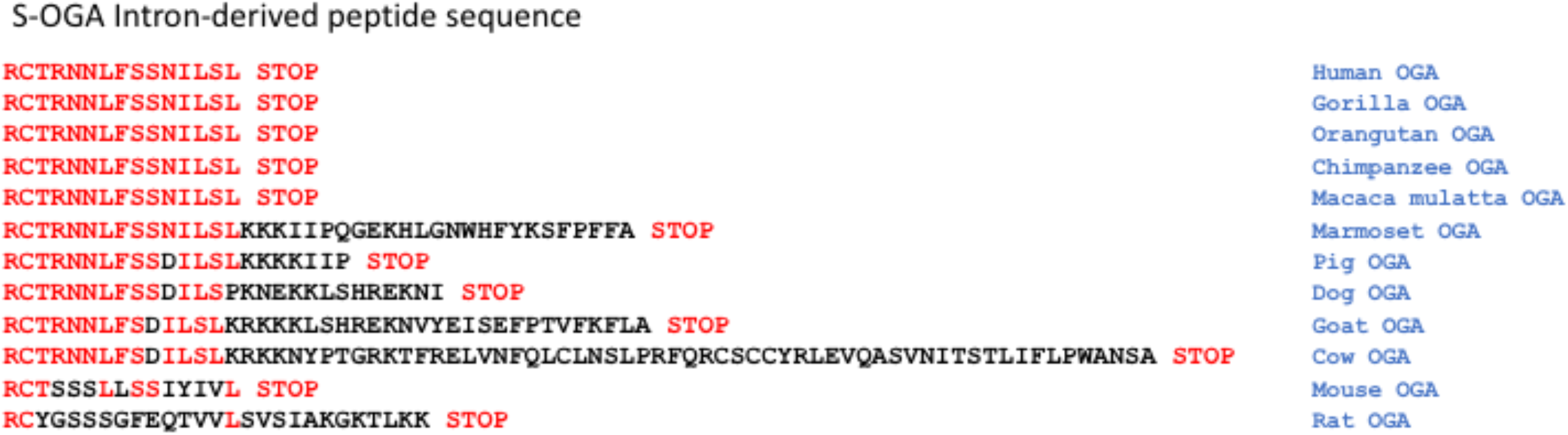
Alignment of the sequences of short OGA intron-derived peptide in different mammalian species.

**Supplementary Figure 4.**
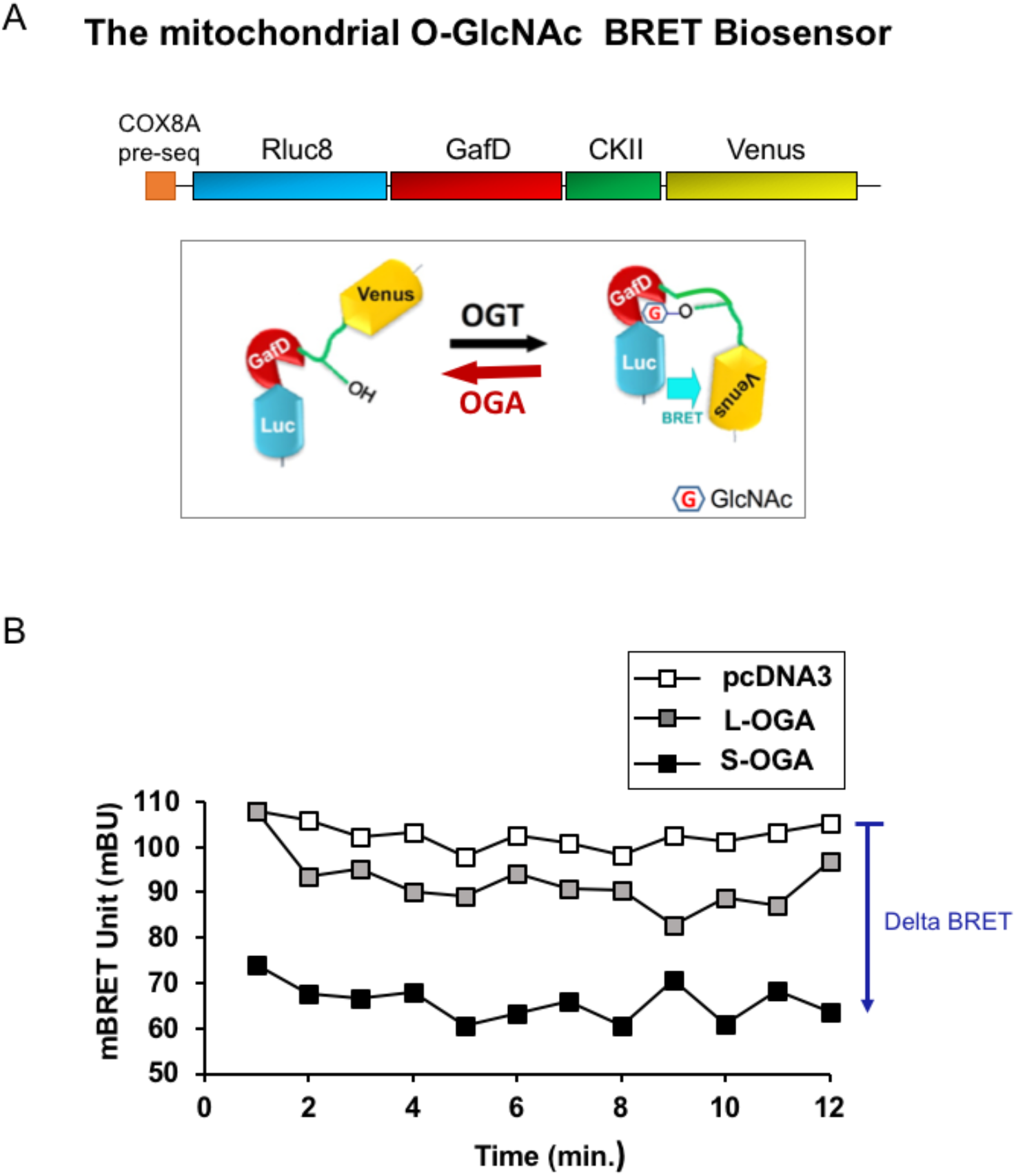
(A) The O-GlcNAc BRET biosensor is composed of Rluc8 luciferase fused to a lectin domain (GafD), a known OGT substrate peptide derived from casein kinase II, followed by the Venus variant of the yellow fluorescent protein. The human COX8A (Cytochrome Oxydase subunit 8A) pre-sequence was inserted upstream of this biosensor for specific targeting to the mitochondria. Basal BRET signal results from the balance between OGT and OGA activities. Upon de-GlcNAcylation by OGA, decreased binding of casein kinase II peptide to GafD lectin domain results in a decrease in BRET signal. (B) Typical experiment showing the monitoring of BRET signal during 10 minutes in pcDNA3, S-OGA and L-OGA transfected HEK-293-T cells. BRET signal was markedly decreased by S-OGA but barely affected by L-OGA.

**Supplementary Table 1:**
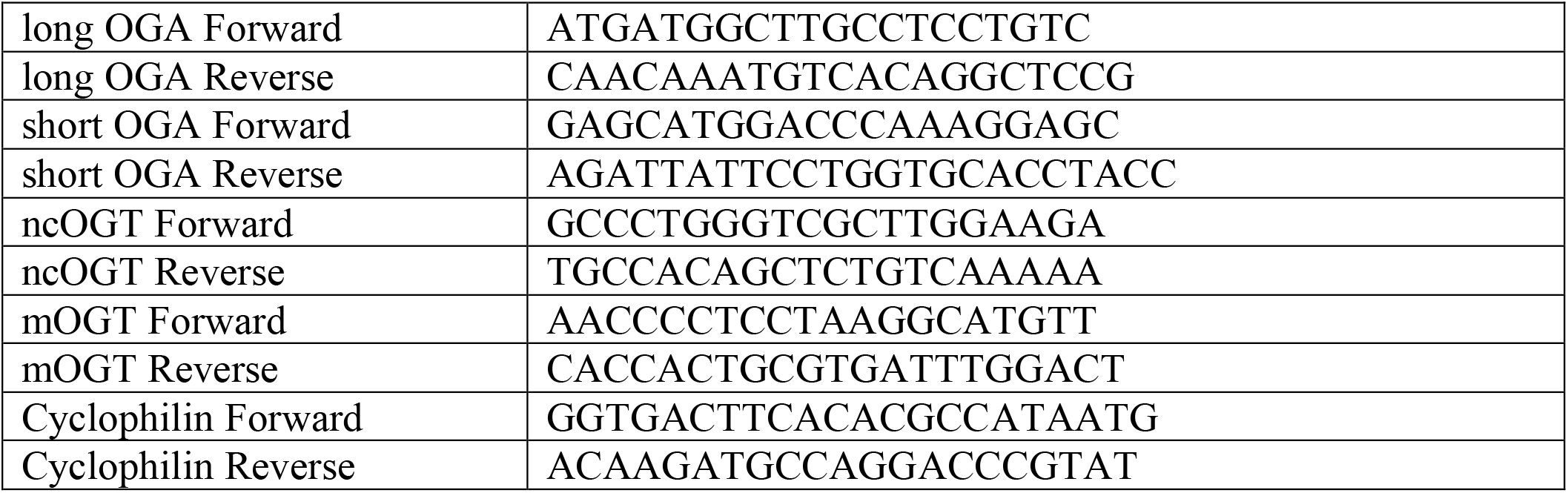
Sequences of oligonucleotides used for RT-qPCR

## References

Akinbiyi EO, Abramowitz LK, Bauer BL, Stoll MSK, Hoppel CL, Hsiao CP, Hanover JA, Mears JA (2021) Blocked O-GlcNAc cycling alters mitochondrial morphology, function, and mass. Sci Rep 11: 22106

Al-Mukh H, Baudoin L, Bouaboud A, Sanchez-Salgado JL, Maraqa N, Khair M, Pagesy P, Bismuth G, Niedergang F, Issad T (2020) Lipopolysaccharide Induces GFAT2 Expression to Promote O-Linked beta-N-Acetylglucosaminylation and Attenuate Inflammation in Macrophages. J Immunol 205: 2499–2510

Banerjee PS, Ma J, Hart GW (2015) Diabetes-associated dysregulation of O-GlcNAcylation in rat cardiac mitochondria. Proc Natl Acad Sci U S A 112: 6050–6055

Blanquart C, Gonzalez-Yanes C, Issad T (2006) Monitoring the activation state of insulin/insulin-like growth factor-1 hybrid receptors using bioluminescence resonance energy transfer. Mol Pharmacol 70: 1802–1811

Bond MR, Hanover JA (2013) O-GlcNAc cycling: a link between metabolism and chronic disease. Annu Rev Nutr 33: 205–229

Candas D, Li JJ (2014) MnSOD in oxidative stress response-potential regulation via mitochondrial protein influx. Antioxid Redox Signal 20: 1599–1617

Chen PH, Chi JT, Boyce M (2018) Functional crosstalk among oxidative stress and O-GlcNAc signaling pathways. Glycobiology 28: 556–564

Comtesse N, Maldener E, Meese E (2001) Identification of a nuclear variant of MGEA5, a cytoplasmic hyaluronidase and a beta-N-acetylglucosaminidase. Biochem Biophys Res Commun 283: 634–640

Gao Y, Wells L, Comer FI, Parker GJ, Hart GW (2001) Dynamic O-glycosylation of nuclear and cytosolic proteins: cloning and characterization of a neutral, cytosolic beta-N-acetylglucosaminidase from human brain. J Biol Chem 276: 9838–9845

Groussaud D, Khair M, Tollenaere AI, Waast L, Kuo MS, Mangeney M, Martella C, Fardini Y, Coste S, Souidi M et al (2017) Hijacking of the O-GlcNAcZYME complex by the HTLV-1 Tax oncoprotein facilitates viral transcription. PLoS Pathog 13: e1006518

Hanover JA, Krause MW, Love DC (2012) Bittersweet memories: linking metabolism to epigenetics through O-GlcNAcylation. Nat Rev Mol Cell Biol 13: 312–321

Hanover JA, Yu S, Lubas WB, Shin SH, Ragano-Caracciola M, Kochran J, Love DC (2003) Mitochondrial and nucleocytoplasmic isoforms of O-linked GlcNAc transferase encoded by a single mammalian gene. Arch Biochem Biophys 409: 287–297

Hu Y, Suarez J, Fricovsky E, Wang H, Scott BT, Trauger SA, Han W, Hu Y, Oyeleye MO, Dillmann WH (2009) Increased enzymatic O-GlcNAcylation of mitochondrial proteins impairs mitochondrial function in cardiac myocytes exposed to high glucose. J Biol Chem 284: 547–555

Issad T, Boute N, Pernet K (2002) A homogenous assay to monitor the activity of the insulin receptor using Bioluminescence Resonance Energy Transfer. Biochem Pharmacol 64: 813–817

Issad T, Kuo M (2008) O-GlcNAc modification of transcription factors, glucose sensing and glucotoxicity. Trends Endocrinol Metab 19: 380–389

Issad T, Masson E, Pagesy P (2010) O-GlcNAc modification, insulin signaling and diabetic complications. Diabetes Metab 36: 423–435

Jozwiak P, Ciesielski P, Zakrzewski PK, Kozal K, Oracz J, Budryn G, Zyzelewicz D, Flament S, Vercoutter-Edouart AS, Bray F et al (2021) Mitochondrial O-GlcNAc Transferase Interacts with and Modifies Many Proteins and Its Up-Regulation Affects Mitochondrial Function and Cellular Energy Homeostasis. Cancers (Basel) 13

Kazemi Z, Chang H, Haserodt S, McKen C, Zachara NE (2010) O-linked beta-N-acetylglucosamine (O-GlcNAc) regulates stress-induced heat shock protein expression in a GSK-3beta-dependent manner. J Biol Chem 285: 39096–39107

Keembiyehetty CN, Krzeslak A, Love DC, Hanover JA (2011) A lipid-droplet-targeted O-GlcNAcase isoform is a key regulator of the proteasome. J Cell Sci 124: 2851–2860

Kim EJ, Kang DO, Love DC, Hanover JA (2006) Enzymatic characterization of O-GlcNAcase isoforms using a fluorogenic GlcNAc substrate. Carbohydr Res 341: 971–982

Lacasa D, Boute N, Issad T (2005) Interaction of the insulin receptor with the receptor-like protein tyrosine phosphatases PTPalpha and PTPepsilon in living cells. Mol Pharmacol 67: 1206–1213

Liu JF, Issad T, Chevet E, Ledoux D, Courty J, Caruelle JP, Barritault D, Crepin M, Bertin B (1998) Fibroblast growth factor-2 has opposite effects on human breast cancer MCF-7 cell growth depending on the activation level of the mitogen-activated protein kinase pathway. Eur J Biochem 258: 271–276

Love DC, Kochan J, Cathey RL, Shin SH, Hanover JA (2003) Mitochondrial and nucleocytoplasmic targeting of O-linked GlcNAc transferase. J Cell Sci 116: 647–654

Ma J, Liu T, Wei AC, Banerjee P, O’Rourke B, Hart GW (2015) O-GlcNAcomic Profiling Identifies Widespread O-Linked beta-N-Acetylglucosamine Modification (O-GlcNAcylation) in Oxidative Phosphorylation System Regulating Cardiac Mitochondrial Function. J Biol Chem 290: 29141–29153

Makino A, Suarez J, Gawlowski T, Han W, Wang H, Scott BT, Dillmann WH (2011) Regulation of mitochondrial morphology and function by O-GlcNAcylation in neonatal cardiac myocytes. Am J Physiol Regul Integr Comp Physiol 300: R1296–1302

Ngoh GA, Watson LJ, Facundo HT, Jones SP (2011) Augmented O-GlcNAc signaling attenuates oxidative stress and calcium overload in cardiomyocytes. Amino Acids 40: 895–911

Nouaille S, Blanquart C, Zilberfarb V, Boute N, Perdereau D, Roix J, Burnol AF, Issad T (2006) Interaction with Grb14 results in site-specific regulation of tyrosine phosphorylation of the insulin receptor. EMBO Rep 7: 512–518

Olenych SG, Claxton NS, Ottenberg GK, Davidson MW (2007) The fluorescent protein color palette. Curr Protoc Cell Biol Chapter 21: Unit 21 25

Oliver DMA, Reddy PH (2019) Small molecules as therapeutic drugs for Alzheimer’s disease. Mol Cell Neurosci 96: 47–62

Pagesy P, Tachet C, Mostefa-Kara A, Larger E, Issad T (2018) Increased OGA expression and activity in leukocytes from patients with diabetes: correlation with inflammation markers Exp Clin Endocrinol Diabetes doi: 10.1055/a-0596-7337.

Pak VV, Ezerina D, Lyublinskaya OG, Pedre B, Tyurin-Kuzmin PA, Mishina NM, Thauvin M, Young D, Wahni K, Martinez Gache SA et al (2020) Ultrasensitive Genetically Encoded Indicator for Hydrogen Peroxide Identifies Roles for the Oxidant in Cell Migration and Mitochondrial Function. Cell Metab 31: 642–653 e646

Sacoman JL, Dagda RY, Burnham-Marusich AR, Dagda RK, Berninsone PM (2017) Mitochondrial O-GlcNAc transferase (mOGT) regulates mitochondrial structure, function and survival in HeLa cells. J Biol Chem, doi: 101074/jbcM116726752 [Epub ahead of print]

Shin SH, Love DC, Hanover JA (2011) Elevated O-GlcNAc-dependent signaling through inducible mOGT expression selectively triggers apoptosis. Amino Acids 40: 885–893

Slawson C, Zachara NE, Vosseller K, Cheung WD, Lane MD, Hart GW (2005) Perturbations in O-linked beta-N-acetylglucosamine protein modification cause severe defects in mitotic progression and cytokinesis. J Biol Chem 280: 32944–32956

Smith RA, Hartley RC, Murphy MP (2011) Mitochondria-targeted small molecule therapeutics and probes. Antioxid Redox Signal 15: 3021–3038

St Amand MM, Bond MR, Riedy J, Comly M, Shiloach J, Hanover JA (2018) A genetic model to study O-GlcNAc cycling in immortalized mouse embryonic fibroblasts. J Biol Chem 293: 13673–13681

Strobel A, Siquier K, Zilberfarb V, Strosberg AD, Issad T (1999) Effect of thiazolidinediones on expression of UCP2 and adipocyte markers in human PAZ6 adipocytes. Diabetologia 42: 527–533

Tan EP, McGreal SR, Graw S, Tessman R, Koppel SJ, Dhakal P, Zhang Z, Machacek M, Zachara NE, Koestler DC et al (2017) Sustained O-GlcNAcylation reprograms mitochondrial function to regulate energy metabolism. J Biol Chem 292: 14940–14962

Tan EP, Villar MT, e L, Lu J, Selfridge JE, Artigues A, Swerdlow RH, Slawson C (2014) Altering O-linked beta-N-acetylglucosamine cycling disrupts mitochondrial function. J Biol Chem 289: 14719–14730

Trapannone R, Mariappa D, Ferenbach AT, van Aalten DM (2016) Nucleocytoplasmic human O-GlcNAc transferase is sufficient for O-GlcNAcylation of mitochondrial proteins. Biochem J 473: 1693–1702

Turkowyd B, Virant D, Endesfelder U (2016) From single molecules to life: microscopy at the nanoscale. Anal Bioanal Chem 408: 6885–6911

Wang X, Feng Z, Wang X, Yang L, Han S, Cao K, Xu J, Zhao L, Zhang Y, Liu J (2016) O-GlcNAcase deficiency suppresses skeletal myogenesis and insulin sensitivity in mice through the modulation of mitochondrial homeostasis. Diabetologia 59: 1287–1296

Zhang Z, Tan EP, VandenHull NJ, Peterson KR, Slawson C (2014) O-GlcNAcase Expression is Sensitive to Changes in O-GlcNAc Homeostasis. Front Endocrinol (Lausanne) 5: 206

Zhao L, Feng Z, Zou X, Cao K, Xu J, Liu J (2014) Aging leads to elevation of O-GlcNAcylation and disruption of mitochondrial homeostasis in retina. Oxid Med Cell Longev 2014: 425705

